# Network-based prediction of protein interactions

**DOI:** 10.1101/275529

**Authors:** István A. Kovács, Katja Luck, Kerstin Spirohn, Yang Wang, Carl Pollis, Sadie Schlabach, Wenting Bian, Dae-Kyum Kim, Nishka Kishore, Tong Hao, Michael A. Calderwood, Marc Vidal, Albert-László Barabási

## Abstract

As biological function emerges through interactions between a cell’s molecular constituents, understanding cellular mechanisms requires us to catalogue all physical interactions between proteins [1–4]. Despite spectacular advances in high-throughput mapping, the number of missing human protein-protein interactions (PPIs) continues to exceed the experimentally documented interactions [5, 6]. Computational tools that exploit structural, sequence or network topology information are increasingly used to fill in the gap, using the patterns of the already known interactome to predict undetected, yet biologically relevant interactions [7–9]. Such network-based link prediction tools rely on the Triadic Closure Principle (TCP) [10–12], stating that two proteins likely interact if they share multiple interaction partners. TCP is rooted in social network analysis, namely the observation that the more common friends two individuals have, the more likely that they know each other [13, 14]. Here, we offer direct empirical evidence across multiple datasets and organisms that, despite its dominant use in biological link prediction, TCP is not valid for most protein pairs. We show that this failure is fundamental - TCP violates both structural constraints and evolutionary processes. This understanding allows us to propose a link prediction principle, consistent with both structural and evolutionary arguments, that predicts yet uncovered protein interactions based on paths of length three (L3). A systematic computational cross-validation shows that the L3 principle significantly outperforms existing link prediction methods. To experimentally test the L3 predictions, we perform both large-scale high-throughput and pairwise tests, finding that the predicted links test positively at the same rate as previously known interactions, suggesting that most (if not all) predicted interactions are real. Combining L3 predictions with experimental tests provided new interaction partners of FAM161A, a protein linked to retinitis pigmentosa, offering novel insights into the molecular mechanisms that lead to the disease. Because L3 is rooted in a fundamental biological principle, we expect it to have a broad applicability, enabling us to better understand the emergence of biological function under both healthy and pathological conditions.

**Summary:** We unveil a fundamental organizing principle of biological networks and demonstrate its predictive power for uncovering novel protein interactions.

---

At the heart of current network-based link prediction algorithms is the triadic closure principle (TCP) [7] (SI Table I.), connecting two nodes if they share multiple neighbors (Fig. 1a). TCP was transplanted into cell biology through the plausible hypothesis that if two proteins have multiple common neighbors, they must participate in the same cellular function, hence they likely also interact with each other [7]. Despite its central role in network-based link prediction algorithms, there is no direct evidence of the validity of the TCP within the subcellular context. To investigate the validity of the TCP hypothesis we measured the number of shared interaction partners of proteins X and Y using the Jaccard similarity *J* = *|NX n NY |/|NX ? NY |*, where *NX* and *NY* are the interaction partners of X and Y. According to the TCP, the higher the Jaccard similarity, the higher is the expected probability that two proteins are connected (Fig. 1b). Strikingly, we find the opposite trend in all PPI networks tested, across different organisms and approaches used to generate interaction datasets (SI A, Table II): The larger the Jaccard similarity between two proteins, the lower the chance that they are known to interact with each other (Figs. 1c, S5). In other words, the starting hypothesis of current network-based PPI prediction tools cannot be validated. This *TCP paradox* is a consequence of taking a well documented pattern in social networks into the biological space, while ignoring the structural and evolutionary mechanisms that govern protein interactions. In the following we propose a biologically sound principle behind the interactome that avoids the pitfalls of TCP and enables us to outperform available link prediction algorithms.

**FIG. 1.**
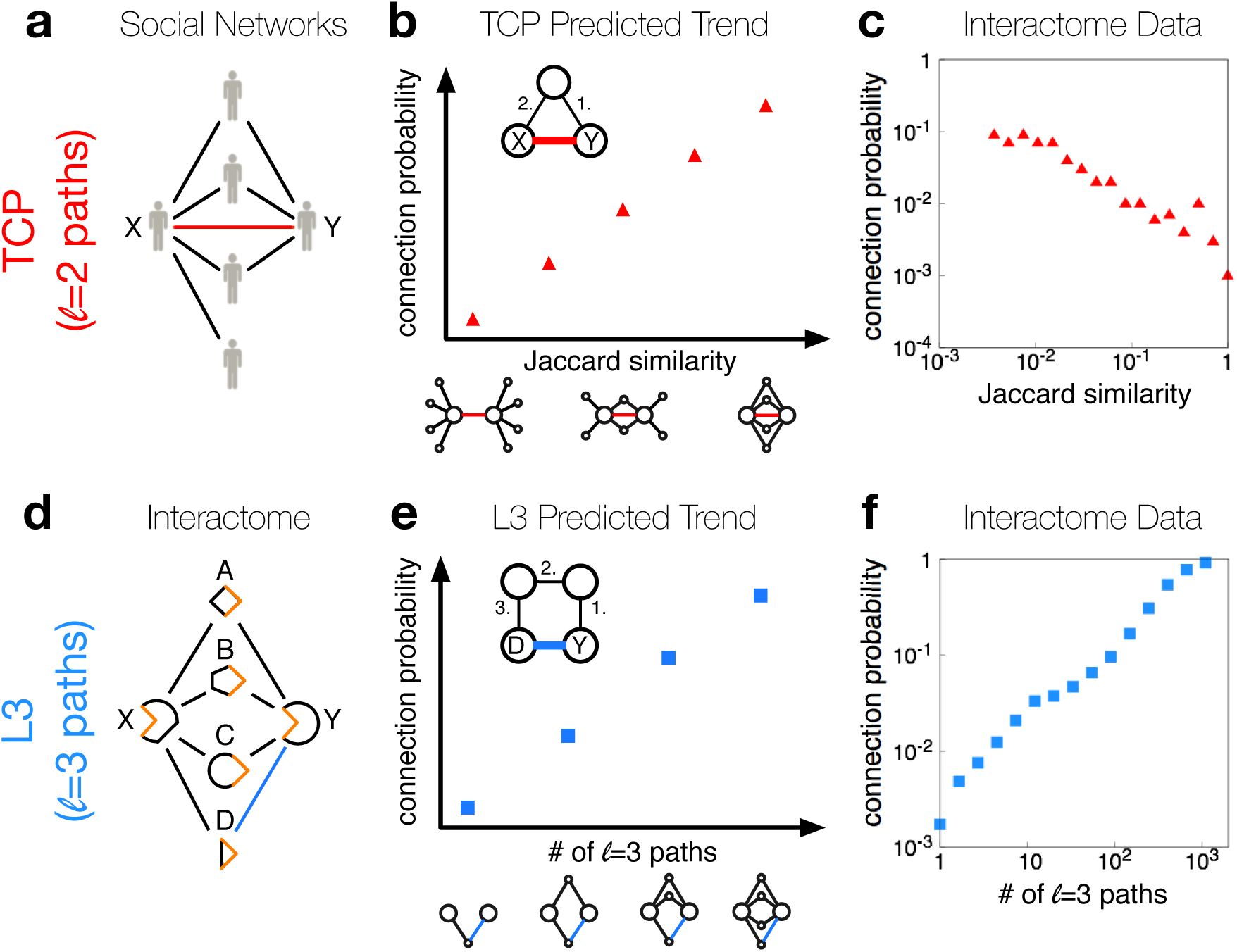
TCP Paradox. (**a**) In social networks, a large number of friends in common implies a higher chance to become friends (red link between nodes X and Y), known as the Triadic Closure Principle (TCP). (**b**) A basic mathematical formulation of TCP states that protein pairs of high Jaccard similarity are more likely to interact. For an alternative similarity measure see Fig. S4.(**c**) We observe the opposite trend in Protein-Protein Interaction (PPI) datasets, as illustrated here for a binary human PPI network (HI-II-14) [20]: high Jaccard similarity indicates a lower chance for the proteins to interact. The data is binned logarithmically based on the Jaccard similarity values. (**d**) PPIs often require complementary interfaces [7, 8], hence two proteins, X and Y, with similar interfaces share many of their neighbors. Yet, X and Y might not directly interact with to interact if linked by multiple *l* = 3 paths in the network (L3). (**f**) As opposed to c), we observe a strong positive trend in HI-II-14 between the probability of two proteins interacting and the number of *l* = 3 paths between them, supporting the validity of the L3 principle.

## Structural Patterns in PPI Networks

According to the TCP hypothesis, if proteins X and Y share multiple interaction partners (A,B,C), they likely interact with each other as well (Fig. 1a). However, from a structural perspective, proteins X and Y are merely expected to share some of their interaction interfaces recognizing the binding sites in proteins A, B, and C (Fig. 1d). Since interaction interfaces of the same type (those in X and Y) do not necessarily interact with each other, there is no reason to expect a direct interaction between X and Y. Instead, if protein X also interacts with protein D, we expect that protein Y is also capable of binding protein D (Fig. 1d). In graph-theoretic terms, TCP predicts that proteins X and Y likely interact with each other if they are connected by paths of length *l*= 2. Instead, the structural argument of Fig. 1d suggests that proteins linked by multiple paths of length *l*= 3, like proteins D and Y, are more likely to have a direct link (L3 principle, Fig. 1e). We arrive to the same conclusion if we rely on evolutionary arguments following gene duplication, predicting again that proteins with multiple shared interaction partners are likely to share even more partners (SI. C) [15–17]. To test the L3 principle, we measured the number of *l*= 3 paths between all protein pairs, finding a direct correlation between the number of *l*= 3 paths between a given protein pair, and the likelihood that they directly interact with each other (Fig. 1f).

*Resolving the TCP paradox:* The structural and evolutionary arguments offer a second testable prediction (Figs. 2a-c, S2): the TCP hypothesis does apply to a small number of interactions, those between self-interacting proteins (SIPs). Indeed, if the interaction domain shared by proteins X and Y can self-interact, potentially forming a homodimer like a SH3-SH3 domain interaction, then X and Y in Fig. 1d are expected to interact, as they share a self-interacting domain. To test this prediction, we consider separately interactions between SIPs and nonSIPs (Fig. 2d-f). We find that, as predicted, the Jaccard similarity between two SIPs correlates with the probability to be linked to each other. The effect is the opposite for PPIs mediated by nonSIPs (Fig. 2d-f), a pattern that holds for a broad range of data sources and organisms (Fig. S6). Given that SIPs follow TCP, why is then TCP violated when all interactions are inspected together (Fig. 2a-c)? The answer is simple: only*∼* 10% of the proteins are known SIPs, and they are altogether involved in only *∼* 2 *-* 4% of the protein pairs (Fig. 2d-f). In other words, TCP, or *l*= 2 connectivity, is expected to fail for more than 96% of potential PPIs. In contrast, the L3 principle applies to all proteins, independent if they are SIPs or nonSIPs (Fig. 2g-i).

**FIG. 2.**
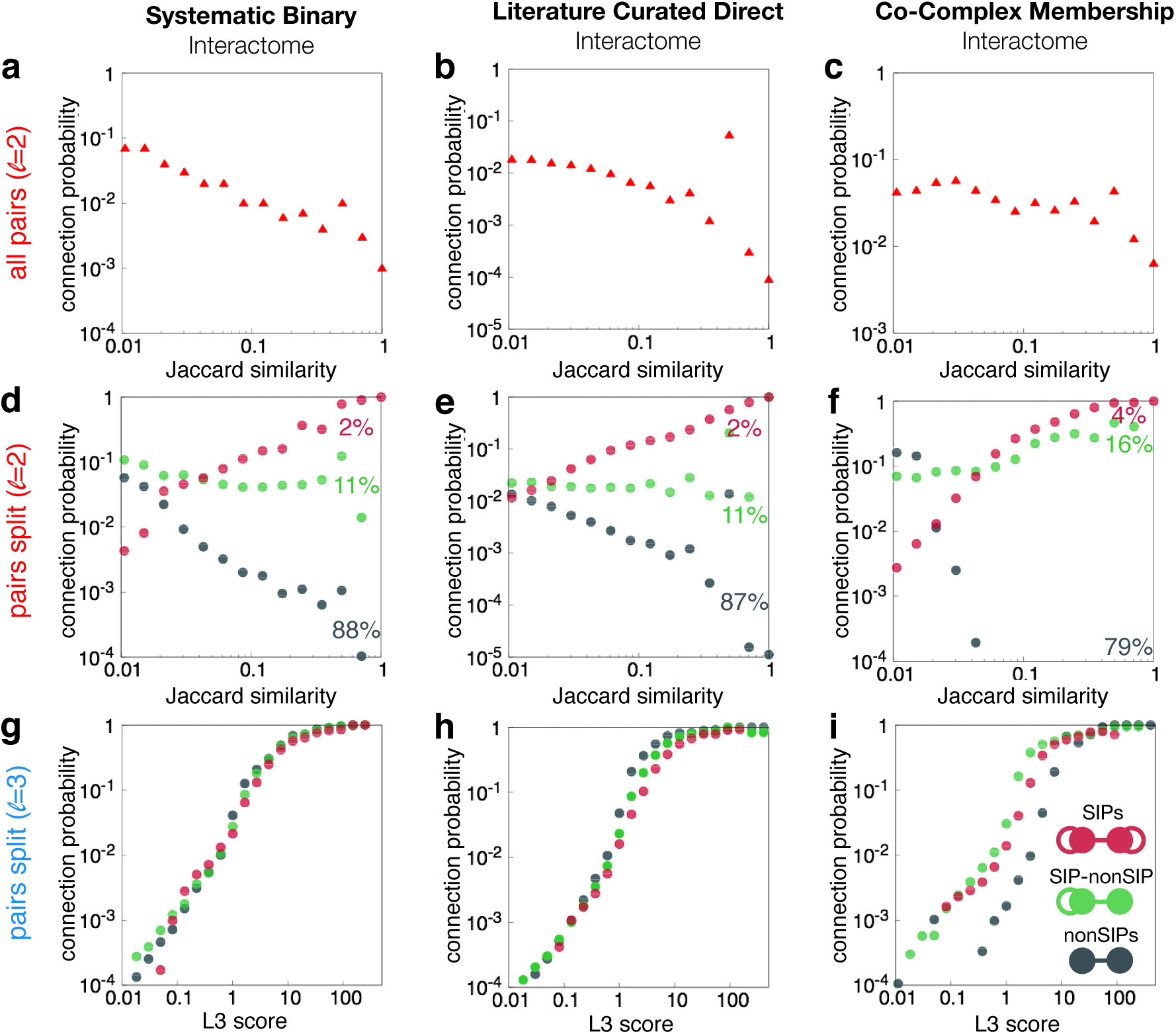
Resolving the TCP paradox. (**a**, **b**, **c**) The lack of positive correlation between accard similarity and connectivity holds for all studied PPI datasets (Fig. S5), as illustrated here for human PPI networks a) obtained by systematic binary methods (HI-II-14 [20]) b) summarizing direct interactions extracted from BioGRID [37] and c) systematically generated using AP-MS techniques, representing co-complex memberships [36]. (**d**, **e**, **f**) Two proteins can either be both self-interacting proteins (SIPs), nonSIPs or a SIP and a nonSIP. We observe positive correlation between Jaccard similarity and connectivity between two SIPs (red); lack of correlation between a SIP and nonSIP (green); and a negative tendency between nonSIPs (grey). Thus, TCP is limited to pairs of SIPs, representing only a small fraction of all protein pairs. (**g**, **h**, **i**) In contrast to Jaccard similarity, *l* = 3 connectivity, defined in Eq. (1), shows a strong positive correlation with the connection probability for all protein pairs. For illustration on additional datasets see Figs. S5 & 6.

*Predicting missing interactions:* We expect that node pairs connected by the highest number of *l*= 3 paths are most likely to be directly connected. However, high degree nodes (hubs) might induce multiple, unspecific shortcuts in the network, biasing the results. To cancel potential degree biases caused by intermediate hubs in the paths, we assign a degree-normalized L3 score to each node pair, *X* and *Y*

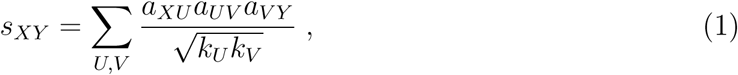

where *aXU* is the interaction strength between nodes X and U and *kU* is the degree of node *U*. Here, as a natural choice offered by graph theory, Eq. (1) measures the connectivity between U and V by the degree-normalized adjacency matrix.

## Computational Cross-validation

To quantify its predictive power, we compare L3 with two basic link prediction algorithms: the Common Neighbors (CN) algorithm [18] (Fig. 3a), a TCP implementation that outperforms Jaccard similarity (Fig. S8) and the Preferential Attachment principle (PA) [18, 19], a link prediction method that mimics unspecific binding between “sticky” proteins by placing a link between two nodes with a score given by the product of their degrees, a method *not* based on TCP. Comparison with other link prediction algorithms is discussed in Figs. S7 and S8. We removed 50% of the PPIs from the interactome maps and typically used the top 500-2000 predicted interactions by each algorithm to determine their precision, measuring the fraction of positive PPIs identified. We find that the L3 algorithm outperforms with a factor of 2 to 3 both CN and PA (Fig. 3a), a substantial improvement that holds for all organisms and data sources tested (Fig. S7 and SI Table III). Interestingly, L3 outperforms all methods even if we consider only SIP pairs, for which TCP is expected to be valid (Fig. S2). For completeness, we checked the performance of paths of lengths up to *l*= 8, finding that the highest precision is indeed provided by *l*= 3 paths (Fig. 3b). Next, we describe two complementary, large-scale experiments to directly test the predictive improvement offered by the L3 principle.

**FIG. 3.**
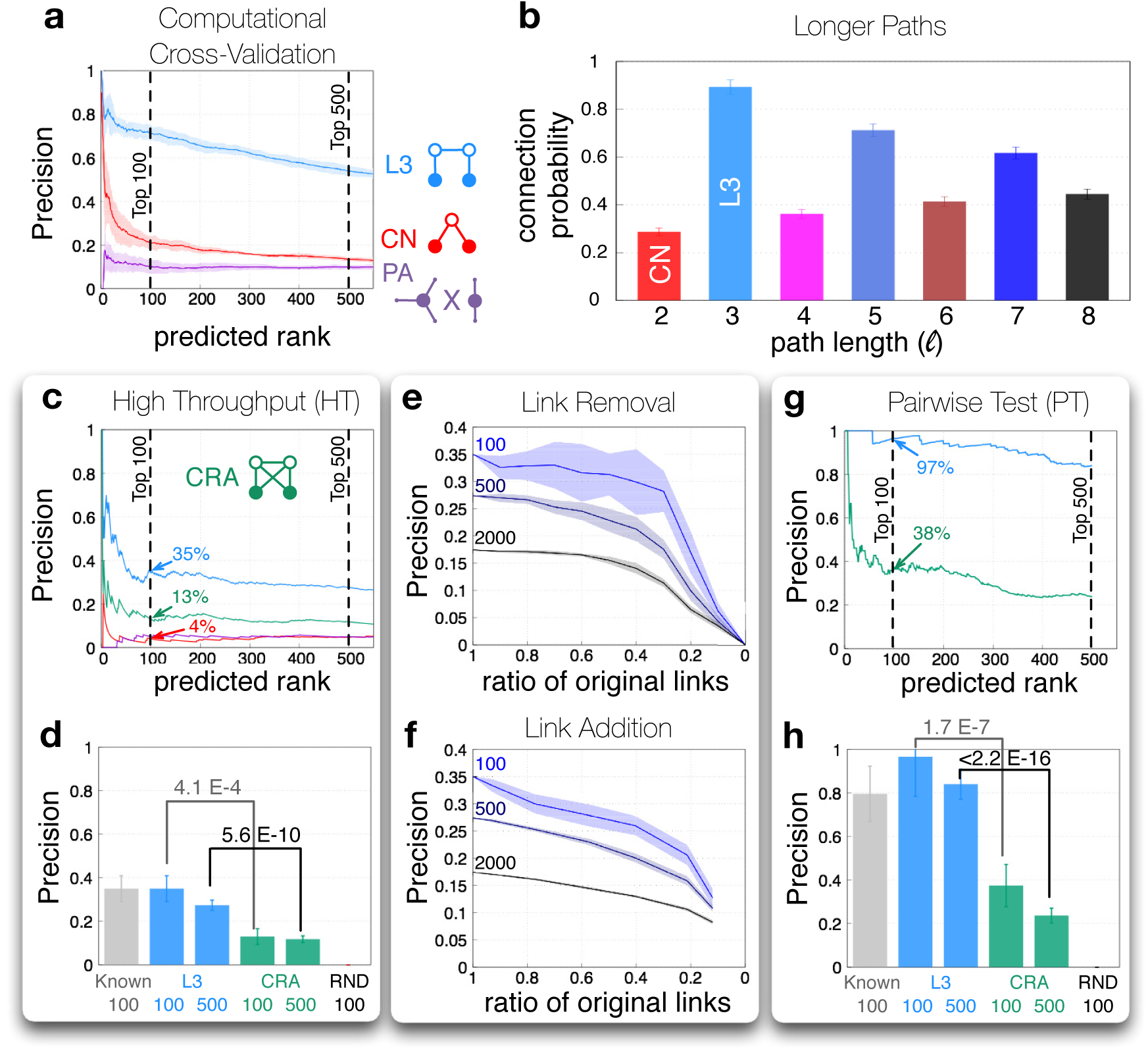
Computational and Experimental Tests. (**a**) Computational cross-validation in HI-II-14 [20]. We find that L3 outperforms CN and PA at least three-fold. The shading around each curve indicates the standard deviation over 10 different random selections, for additional datasets see Fig. S7. (**b**) Connection probability in the top 1,000 HI-II-14 protein pairs ranked by different powers of the adjacency matrix, *l*, counting all paths of length *l* = 2, …, 8. *l* = 3 paths are the most informative on direct connectivity. (**c**) In a high-throughput (HT) setting, we tested the L3 predictions on the human interactome HI-filtered, against the HT human interactome, HI-III [21]. CRA was found to be the best performing algorithm out of 23 different methods tested (Fig. S8). As a positive benchmark, we selected 100 known HI-filtered interactions (“Known”) and as a negative benchmark, 100 random pairs (“RND”), for details see SI B. (**d**) The recovery rate (precision) of L3 is significantly higher than that of CRA and comparable to the one of known interactions.(**e**) Robustness analyses of the L3 predictions with HT validation against data incompleteness, evaluated at the top 100, 500 and 2000 predictions, respectively. L3 is robust even when less then half of the PPIs are kept. (**f**) L3 is also robust against adding random links to the network, even when less then half of the links are PPIs. (**g**) Pairwise testing the top 500 predictions of L3 and CRA. We indicate the pairs where the experiments were conclusive (positive or negative) (SI B).(**h**) L3 not only outperforms CRA, but the L3 predictions test positively with about the same rate as known interactions, indicating an optimal performance.

## High-Throughput Experiments

To enable a precise evaluation, we used a filtered version of the human interaction dataset HI-II-14 [20], (HI-filtered, SI. A & B) to predict new PPIs using Eq. (1). We test our predictions using a new human binary PPI map, HI-III [21], generated by screening a search space of *∼* 18, 000 *×* 18, 000 human genes for PPIs that entirely contains the search space of HI-filtered. Since HI-III is an independently generated, still incomplete, high-throughput (HT) dataset, we do not expect to recover all interacting pairs. Can we estimate the best possible recovery rate for the L3 predictions? To do so, we selected as positive control 100 known interactions in HI-filtered, involving at least one of the proteins in the top 500 L3 predictions. We find that 35% of these established interactions are recovered in the HT test. This rate is somewhat lower than the 42% recovery rate observed on average for all HI-filtered interactions, indicating that the tested proteins might be somewhat harder to address experimentally. Altogether, a 35% recovery rate of the predicted interactions would indicate that most (if not all) of them are true interactions. As negative control, we randomly selected 100 non-interacting pairs, involving at least one of the proteins in the top 500 L3 predictions (“RND”). We did not recover any of these pairs in HI-III (Fig 3f, SI B). When testing the predicted interactions, L3 significantly outperforms not only CN and PA (Fig. 3a), but also each of the 21 other network-based link prediction methods tested (Fig. S8). Remarkably, the top 100 L3 predictions have a 35% recovery rate, the maximal possible value. This precision is seven fold higher than either CN or PA and more than twice the precision of the best performing literature method, Community Resource Allocation (CRA) [22] (Fig. 3c-d). CRA outperforms other TCP-based methods because it counts network motifs containing both *l*= 2 and *l*= 3 paths (Fig. 3c), offering a subset of the L3 predictions. Yet, CRA’s inferior performance to L3 is rooted in its reliance on *l*= 2 connectivity as well, that acts as a biased decimation. We also tested if our results are robust against data incompleteness, finding that the precision of the predicted interactions is stable up to removal of even 60 *-* 70% of the known interactions (false negatives, Fig. 3e), or when the number of randomly added links exceeds the number of original links (false positives, Fig. 3f).

## Pairwise Tests

HI-III underestimates the performance of L3 as only a fraction of the existing PPIs are detectable in a HT setting [5]. We therefore also assessed the performance of L3 in yeast-two-hybrid pairwise tests (PT) for the top 500 predicted links, utilizing – amongst others – the same positive and negative controls as above (Fig. 3g, SI B). Altogether, we performed *∼* 3, 000 pairwise tests, classifying each pair as either positive, negative or undetermined. The recovery rate of the known interactions – i.e. the fraction of positives over positives and negatives – has now increased to 80%, while none of the negative control pairs scored positive (Fig. 3h). The recovery rate of L3 is once again three fold higher than that of CRA and it is comparable to the positive control, i.e. 84% for the top 500 L3 predictions (Fig. 3h). These results can be computationally extrapolated to larger scales and further datasets, using a leave-one-out cross-validation approach (Fig. S11). Thereby, we estimate that *∼* 6, 000 of the top 10, 000 L3 predictions are true PPIs when using HI-III to predict new PPIs, a yield exceeding the *∼* 4 *-* 5, 000 novel interactions provided by an additional HT screen (HI-III [21]). This indicates that L3 could be an accurate guide for future high-throughput testing, helping prioritize the search space.

## Retinitis Pigmentosa

Previously undetected PPIs can offer novel insights into disease mechanisms. Indeed, our top L3 prediction is an interaction between FAM161A and GOLGA2 (Fig. 4), which was identified in HI-III and tested positively in our pairwise experiment as well. Family-based studies and homozygosity mapping have linked coding mutations in the gene FAM161A to a hereditary form of Retinitis Pigmentosa (RP), a retinal ciliopathy leading to progressive degeneration of photoreceptors. Yet, the cellular functions of FAM161A as well as the molecular mechanisms leading to RP upon loss of FAM161A are largely unknown [23]. FAM161A localizes at the ciliary basal body and the connecting cilium of human photoreceptors [24, 25], binds to microtubules [25] and has recently been found to interact with proteins of the Golgi apparatus (GA) [26]. The predicted and confirmed interaction between FAM161A and GOLGA2, a core member of the GA, offers a novel mechanistic insight for the role of FAM161A in GA function. Remarkably, FAM161A and GOLGA2 do not share any interaction partners, hence TCP-based algorithms are unlikely to predict this interaction. Within the top 500 L3 predictions we find five additional experimentally confirmed interaction partners of FAM161A: TRIM23, VPS52, KIFC3, TRAF2, and REL (Fig. 4), each of them offering insights into the function of FAM161A. TRIM23 and VPS52 both show GA localization and have been implicated in regulating vesicle trafficking between the GA and other cellular compartments. KIFC3, a recently identified interactor of FAM161A [26], is well known for its function as a microtubule motor protein. Our findings can also indicate novel directions, such as a potential link of FAM161A to NFk-b signaling. Indeed, TRIM23, TRAF2 and REL have demonstrated roles in NFk-b activation, which is particularly interesting in light of the activation of cell death pathways in RP-affected photoreceptors. Altogether, these newly identified interactions position FAM161A within a molecular network that connects GA function with cilium organization and intracellular transport, providing detailed insights into potential molecular mechanisms that underlie RP.

**FIG. 4.**
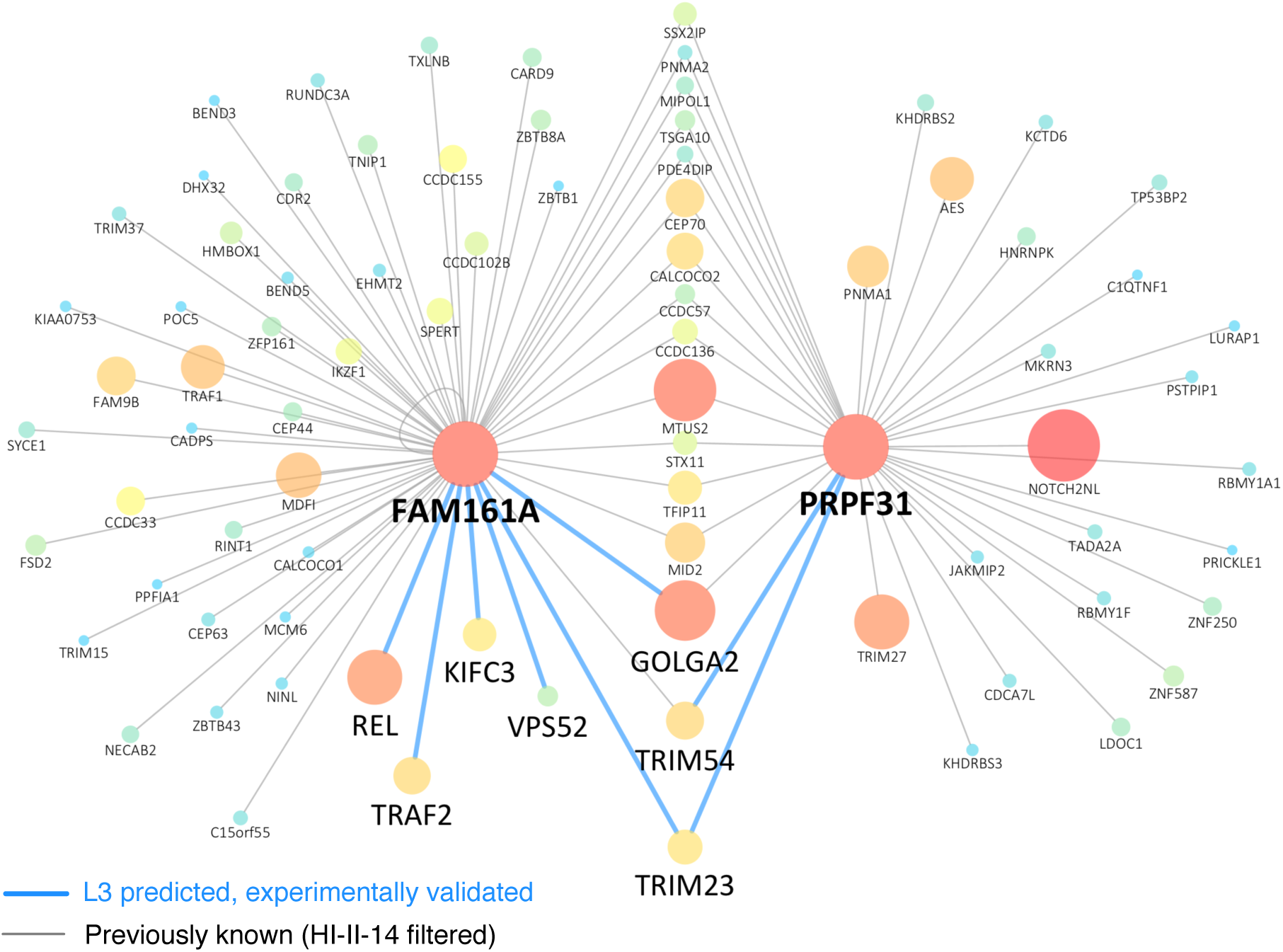
Retinitis Pigmentosa. For two proteins involved in retinitis pigmentosa, FAM161A and PRPF31, we show all known interacting partners in HI-filtered (grey), together with those predicted by the L3 algorithm and confirmed by pairwise tests (blue). The top L3 predicted interaction is connecting FAM161A to GOLGA2, two proteins without any shared interaction partners, beyond the scope of TCP-based methods. The node size and color illustrates the degree of the proteins in HI-filtered. In light of our experiments, GOLGA2, TRIM23, and TRIM54 are now amongst several shared interaction partners between FAM161A and PRPF31, a pre-mRNA splicing factor, whose mutations are causal for another form of RP [23]. This illustrates the key principle behind L3 (Fig. 1d), that two proteins, like FAM161A and PRPF31, despite sharing multiple interacting partners, do not necesseraly interact with each other, but share additional, previously unrecognized interaction partners.

## Discussion

Protein-protein interactions (PPIs) play a central role in the mechanistic understanding of cellular function under normal and disease conditions [27–31]. However, the vast number of missing interactions limit our ability to fully exploit the predictive and explanatory power of the interactome. Existing tools for protein interaction prediction using network topology information are based on principles rooted in social networks. We showed that these principles mechanistically fail for most protein pairs. Instead, we demonstrate and experimentally confirm the predictive power of *l*= 3 paths, a mechanism supported by both structural and evolutionary arguments.

The L3 framework, despite its exceptional predictive power, is not without limitations. First, like all network-based methods, it cannot find interacting partners for proteins without known links. For such proteins, we can, in principle, seamlessly integrate into the L3 framework information on sequence, evolutionary history or 3D structure, used by some PPI prediction algorithms [7, 8, 32–34]. Additionally, sampling biases of the available network datasets can cause false positives. This applies to literature curated PPI networks, where the less explored proteins lack disproportionally more links than disease-related, highly studied hubs, like P53. It also applies to networks obtained by mass-spectrometry [35, 36], where the observed degree of a bait is closer to the degree in the complete network, exceeding the degrees of proteins not yet used as baits. Despite these limitations, we believe that L3 coupled with experimental validation can help us exploit the promise of the interactome in unveiling biological mechanisms.

## ACKNOWLEDGMENTS

We thank D. Hill, F. Roth, F. Cheng and M. Santolini for useful discussions on the manuscript. This work was supported by an NHGRI Center of Excellence in Genome Science grant P50HG004233. We gratefully acknowledge the support of The National Human Genome Research Institute (NHGRI) of NIH; The Ellison Foundation, Boston, MA; and The Dana-Farber Cancer Institute Strategic Initiative.

**Code availibality:** The code written for and used in this study is available from the corresponding author upon reasonable request.

**Data availability:** The human interactome, HI-123 is available at Ref. 21. Additional data that support the findings of this study are available within the supplementary information files or from the corresponding author upon reasonable request.

## AUTHOR CONTRIBUTIONS

Computational analyses were developed and performed by I.A.K., Experiments were performed by K.S., C.P., S.S., D.-K.K., N.K., W.B. processed Sanger sequencing data and T.H. designed the cherrypicking files. Y. W. carried out the protein structural analyses. Principal investigators overseeing primary data management, structural biology and experiment evaluation were A.-L.B., M.V. and M.A.C., A.-L.B. advised the overall research effort. I.A.K., K.L. and A.-L.B. wrote the manuscript with contributions from other co-authors. The authors declare no competing financial interest.

